# A thorough RNA-seq characterization of the porcine sperm transcriptome and its seasonal changes

**DOI:** 10.1101/506899

**Authors:** M. Gòdia, M. Estill, A. Castelló, S. Balasch, J.E. Rodríguez-Gil, S.A. Krawetz, A. Sánchez, A. Clop

**Affiliations:** Animal Genomics Group, Centre for Research in Agricultural Genomics (CRAG) CSIC-IRTA-UAB-UB, Campus UAB, Cerdanyola del Vallès (Barcelona), Catalonia, Spain; Department of Obstetrics and Gynecology, Wayne State University, Detroit, Michigan, USA.; Unit of Animal Science, Department of Animal Science and Nutrition, Autonomous University of Barcelona, Cerdanyola del Vallès (Barcelona), Catalonia, Spain; Grup Gepork S.A. El Macià, Masies de Roda (Barcelona), Catalonia, Spain; Unit of Animal Reproduction, Department of Animal Medicine and Surgery, Autonomous University of Barcelona, Cerdanyola del Vallès (Barcelona), Catalonia, Spain; Center for Molecular Medicine and Genetics, Wayne State University, Detroit, Michigan, USA.; C.S. Mott Center for Human Growth and Development, Wayne State University, Detroit, Michigan, USA.; Consejo Superior de Investigaciones Científicas (CSIC), Barcelona, Catalonia, Spain.

**Keywords:** sperm, sperm RNA element, RNA-seq, sperm seasonality, transcript integrity, differential gene expression

## Abstract

Understanding the molecular basis of cell function and ultimate phenotypes is crucial for the development of biological markers. With this aim, several RNA-seq studies have been devoted to characterize the transcriptome of ejaculated spermatozoa in relation to sperm quality and fertility. Semen quality follows a seasonal pattern and decays in the summer months in several animal species. The aim of this study was to deeply profile the transcriptome of the boar sperm and to evaluate its seasonal changes. We sequenced the total and the short fractions of the sperm RNA from 10 Pietrain boars, 5 collected in summer and 5 five sampled in winter, and identified a complex and rich transcriptome with 4,436 coding genes of moderate to high abundance. Transcript fragmentation was high but less obvious in genes related to spermatogenesis, chromatin compaction and fertility. Short non-coding RNAs mostly included piwi-interacting RNAs, transfer RNAs and micro-RNAs. We also compared the transcriptome of the summer and the winter ejaculates and identified 34 coding genes and 7 micro-RNAs with a significantly distinct distribution. These genes were mostly related to oxidative stress, DNA damage and autophagy. This is the deepest characterization of the boar sperm transcriptome and the first study linking the transcriptome and the seasonal variability of semen quality in animals. The annotation described here can be used as a reference for the identification of markers of sperm quality in pigs.

## Introduction

### Semen quality is highly relevant for the sustainability of modern pig breeding

Swine, together with poultry, are the most important sources of meat for human consumption (in kg) worldwide (OECD, 2018). Moreover, the global demand for animal protein is growing quickly. Thus, improving the efficiency of pork production is of paramount importance for the sustainability of the sector. Pig production relies on the genetic merit of boars kept in artificial insemination centers and the quality of their sperm to disseminate their genetic material. Hence, there is an increasing demand for molecular markers that afford early prediction of semen quality and fertility in young boars.

### The sperm cell contains a complex and functionally relevant transcriptome

For decades, the ejaculated mature sperm was considered a dormant cell that only carried the paternal genome to the egg. Nonetheless, in the recent years the biological complexity of sperm has become more evident, with the discovery of a rich sperm RNA population with functional roles in spermatogenesis, fertilization, early embryo development and transgenerational epigenetic transmission (Gòdia et al., 2018b). Mature sperm RNAs have been studied by Next Generation Sequencing (NGS) in several mammalian species including human (Sendler et al., 2013), horse (Das et al., 2013), mouse (Johnson et al., 2015) and cattle (Selvaraju et al., 2017). These studies have shown a sperm-specific transcriptome with a large population of transcripts most of which are present at low levels and are also highly fragmented. The small non-coding RNA (sncRNA) population of sperm has also been interrogated in several mammals (Krawetz et al., 2011; Das et al., 2013; Capra et al., 2017), and is composed of a large and complex repertoire of microRNAs (miRNAs), piwi-interacting RNAs (piRNAs) and transfer RNAs (tRNAs), among other RNA classes. The abundance of these transcripts has been proposed as a valuable source of bio-markers for semen quality in animal breeding and bio-medicine (Jodar et al., 2015; Salas-Huetos et al., 2015; Capra et al., 2017).

### The boar sperm transcriptome

The boar sperm transcriptome has been interrogated in several studies, most employing qPCR analysis of target genes. Although qPCR is a useful tool that provides very valuable information, these studies typically assume transcript integrity and target one or two exons of only candidate genes. RNA-seq overcomes these two limitations. The first genome wide evaluation of the boar spermatozoa transcriptome was completed in 2009 by sequencing the 5’-ends of a Expressed Sequence Tag library using Sanger technology (Yang et al., 2009), which led to the identification of 514 unique sequences many of which corresponded to unknown genes. High-throughput RNA-seq was more recently applied to compare two differentially fed boars (Bruggmann et al., 2013) and to explore the short RNA component of the boar sperm (Luo et al., 2015; Pantano et al., 2015; Chen et al., 2017a; Chen et al., 2017b). These studies aimed to compare the sncRNAs at different stages of spermatogenesis or between the different components of the ejaculate, and concluded that a large proportion of these short RNAs are sperm-specific. Despite these previous studies, an in-depth analysis of the boar sperm transcriptome is still missing.

### Sperm quality has a seasonal component

Sperm quality can be influenced by multi-factorial genetics (Marques et al., 2017) and environmental factors such as stress and seasonality (Wettemann et al., 1976). In pigs, a clear drop on semen quality and male fertility has been observed in the warm summer months, possibly due to heat stress (Trudeau and Sanford, 1986; Zasiadczyk et al., 2015). This seasonal effect has been linked to altered levels of some transcripts (Yang et al., 2010).

The first step towards the efficient identification of RNA markers of sperm quality requires obtaining a profound picture of the boar sperm transcriptome. Our group has recently optimized a pipeline to extract RNA from swine mature spermatozoa and obtain a high quality and complete transcriptome profile (Gòdia et al., 2018a). In this study, we have profiled the sperm transcriptome from 10 boars, including both coding and non-coding RNAs and we have evaluated the relationship between transcript abundance and the season of collection (summer versus winter) in the northern temperate climate zone.

## Materials and Methods

### Sample collection

Specialized professionals obtained fresh ejaculates from 10 Pietrain boars from a commercial farm, with ages ranging from 9 to 28 months old, between July 2015 and January 2017 as previously described (Gòdia et al., 2018a). Of the 10 ejaculates, 5 were collected between December to February, and the other between May and July. No animal experiment has been performed in the scope of this research.

### RNA extraction, qPCR validation, library prep and sequencing

RNA extraction was performed and the abundances of the sperm specific *PRM1* and the somatic-cell specific *PTPRC* transcripts as well as the presence of genomic DNA (gDNA) were measured by qPCR to determine the quality of the obtained RNAs as previously described by our group (Gòdia et al., 2018a). Extracted RNA was quantified with QubitTM RNA HS Assay kit (Invitrogen; Carlsbad, USA) and its integrity validated with Bioanalyzer Agilent RNA 6000 Pico kit (Agilent Technologies; Santa Clara, USA). Total RNA was subjected to ribosomal RNA depletion with the Ribo-Zero Gold rRNA Removal Kit (Illumina) and RNA-seq libraries were constructed with the SMARTer Low Input Library prep kit v2 (Clontech) and sequenced to generate 75 bp paired-end reads in an Illumina’s HiSeq2500 sequencing system. Short RNA-seq libraries were prepared from the same RNA aliquots (prior to rRNA depletion) with the NEBNext Small RNA (New England Biolabs) and sequenced in an Illumina Hiseq2000 to produce 50 bp single reads.

### Total RNA-seq mapping and analysis of the Sperm RNA Elements

The quality of the paired end reads were evaluated with FastQC v.0.11.1 (http://www.bioinformatics.bbsrc.ac.uk/projects/fastqc), and filtered to remove low quality reads and adaptors with Trimmomatic v.0.36 (Bolger et al., 2014). Filtered reads were then mapped to the *S.scrofa* genome (Sscrofa11.1) with HISAT2 v.2.1.0 (Kim et al., 2015) with default parameters except “--max seeds 30” and “-k 2”. Duplicate mapped reads were removed using Picard Tools (http://picard.sourceforge.net) MarkDuplicates. The uniquely mapped reads were used for the detection and quantification of Sperm RNA Elements (SREs). SREs are short-size sequences characterized by a number of RNA-seq reads clustering to a given genomic location (Jodar et al., 2015; Gòdia et al., 2018b). This approach enables an accurate exon-quantification (or short-size sequence quantification) instead of a whole transcript mean, which makes it useful for highly degraded tissues such as sperm. After mapping, SREs are classified as exonic (mapping to annotated exons), intronic, upstream/downstream 10 kb (if located 10 kb upstream or downstream of annotated genes) and orphan (mapping elsewhere in the genome) (Gòdia et al., 2018b). This classification was done using the pig Ensembl genome annotation (v.91) extracted with the R package “BiomaRt” (Durinck et al., 2009). Porcine orphan SREs coordinates were converted to human (hg38) coordinates and from human to bovine (bosTau8) using the UCSC liftover tool (Kuhn et al., 2013). The coefficient of variation (CV) of the RNA abundance across samples was used to classify the transcripts as highly stable (CV > 0.75), moderately stable (CV between 0.25 and 0.75) and highly unstable (CV < 0.25). Only these genes with all their SREs fitting the same stability class were considered for the GO analysis.

### *De novo* transcriptome analysis

Reads unmapped to the Sscorfa11.1 genome were screened against the porcine Transposable Elements from the Repbase database (Bao et al., 2015) using HISAT2 v.2.1.0 (Kim et al., 2015). The remaining unmatched reads were searched against bacterial and viral genomes using Kraken v.0.10.5 (Wood and Salzberg, 2014) and removed. The remaining reads were subjected to *de novo* assembly with Trinity v.2.1.0 (Grabherr et al., 2011) using default parameters and databases. The assembled contigs were quantified with RSEM and only those with identity score > 85%, abundance levels > 50 FPKM and detected in 5 samples or more were kept.

### Repetitive Elements and long non-coding RNAs

The proportion of reads in Repeat Elements (RE) was calculated with Bedtools (Quinlan and Hall, 2010) multicov using the RepeatMasker database (Bao et al., 2015). Read counts were normalized for RE length and sequencing depth. The same approach was used for long non-coding-RNAs (lncRNAs). Only the lncRNAs annotated in Ensembl v.91 were used. The coding genes mapping less than 20 kb apart from the lncRNAs were considered as potential cis-regulated lncRNA targets.

### Transcript Integrity

RNA transcript integrity (TIN) was calculated with RseQC v.2.6.4 (Wang et al., 2012) using the Ensembl v.91 pig annotation. TIN indicates the proportion of a gene that is covered by reads. As an example, TIN = 100 indicates a fully covered transcript. Transcript abundance was calculated using expression.py from the same software. Transcript length was calculated based on CDS length, extracted with the R package “BiomaRt” (Durinck et al., 2009).

### Analysis of the short non-coding RNAs

Trimming of adaptors and low quality bases were performed with Cutadapt v1.0 (Martin, 2011) and evaluated with FastQC v.0.11.1 (http://www.bioinformatics.bbsrc.ac.uk/projects/fastqc). The mapping of sncRNAs was performed with the sRNAtoolbox v.6.17 (Rueda et al., 2015) with default settings and giving as library datasets: tRNA database (Chan and Lowe, 2016), miRBase (Kozomara and Griffiths-Jones, 2011) release 21, piRNA database (Rosenkranz, 2016) and Mt tRNA, Mt rRNA, snRNA, snoRNA, lincRNA, CDS and ncRNAs from Ensembl v.91. Multi-adjusted read counts were then normalized by sequencing depth. We only considered the miRNAs that were detected in all the samples processed. To determine if piRNAs were located in RE, the overlap between REs and the piRNA clusters that were shared in at least 3 samples was checked with Bedtools (Quinlan and Hall, 2010) multicov using the RepeatMasker database (Bao et al., 2015). The short RNA-seq reads that did not align to any of the datasets provided were used for the *de novo* piRNA annotation using ProTRAC v.2.4.0 (Rosenkranz and Zischler, 2012) and forcing a piRNA length between 26 and 33 bp and a default minimum cluster length of 5 kb. We then kept only these putative novel clusters that were shared in at least 3 of the sperm samples.

### Analysis of the seasonal variation of the boar sperm transcriptome

We studied the potential seasonal effect of the sperm trancriptome by comparing the summer (N=5) and the winter (N=5) ejaculates. Total RNA-seq analysis was performed for the transcripts annotated in the pig genome. We quantified RNA abundance with the software StringTie (Pertea et al., 2015). Transcript counts were then used for the differential analysis after correcting for the sequencing run using the R package DESeq2 (Love et al., 2014) correcting for sequencing run batch. Similarly, the identification of differential miRNAs was also carried with DESeq2 (Love et al., 2014). We only considered the differentially abundant transcripts and miRNAs with adjusted FDR values < 0.05 and FC > 1.5. Gene Ontology enrichment was performed with Cytoscape v.2.3.0 plugin ClueGO v.2.3.5 (Bindea et al., 2009) with the porcine dataset and default settings, only significant corrected p-values with Bonferroni were considered.

## Results and Discussion

### Total RNA-seq analysis: characterization of sperm RNA elements

RNA extraction yielded an average of 2.1 fg per cell (Supplementary File 1). These RNAs were devoid of intact ribosomal 18S and 28S RNA and were free of gDNA and RNA from somatic cell origin [22]. On average, the total RNA-seq libraries yielded 29.5 M paired-end reads (Supplementary File 1). A total of 81.3% of the reads that passed the quality control filter mapped unambiguously to the pig genome (Supplementary File 1). After duplicate removal, a mean of 5.6 M reads per sample were obtained, resulting in a percentage of unique reads similar to recent data on human sperm (unpublished results). These reads were used for further analysis and yielded 185,037 SREs (see the Methods section) (Gòdia et al., 2018b). Most SREs were present at low abundances but the 10% most abundant (top decile) SREs accounted for 65% of the read count with RNA levels ranging between 83 and 378,512 RPKM (Figure 1). Most of these top decile SREs were exonic (Supplementary File 2). Notably, the majority (65%) of the intronic and upstream/downstream 10 kb SREs mapped in or near genes that also harbored exonic SREs. The exonic, intronic and the upstream/downstream 10 kb top decile SREs (see the Methods) mapped in or near 4,436 annotated genes, which were thus considered to be abundant in the boar sperm transcriptome (Supplementary File 2). The top decile SREs also included 2,667 orphan SREs (SREs located more than 10 kb apart from the closest annotated gene) (Supplementary File 2). However, nearly 30% of the orphan SREs mapped within 30 kb from the closest gene, which indicates that, as the novel upstream/downstream 10 kb SREs, they may represent unannotated exons of these genes. In summary, only 10% of the top decile SREs were not linked to annotated genes. A recent study carried by Pertea *et al*. (Pertea et al., 2018) analyzed RNA-seq data from 9,795 human experiments from the GTEx project and concluded that the human genome annotation incorporates most of the *Homo sapiens* genes but still lacks a large proportion of the splice isoforms. While this study increased the list of coding genes by only 5%, the catalogue of splice isoforms grew by 30%. Our data is in line with these recent results and does not only indicate that the novel annotation of the pig genome annotation incorporates most of the genes found in sperm but also reveals that there is still a large amount of splice isoforms to be discovered in this species. Since it is well known that the spermatozoon harbors a very specific transcriptome, a large proportion of these unannotated isoforms are likely to be sperm-specific (Sendler et al., 2013; Ma et al., 2014).

**Figure 1.**
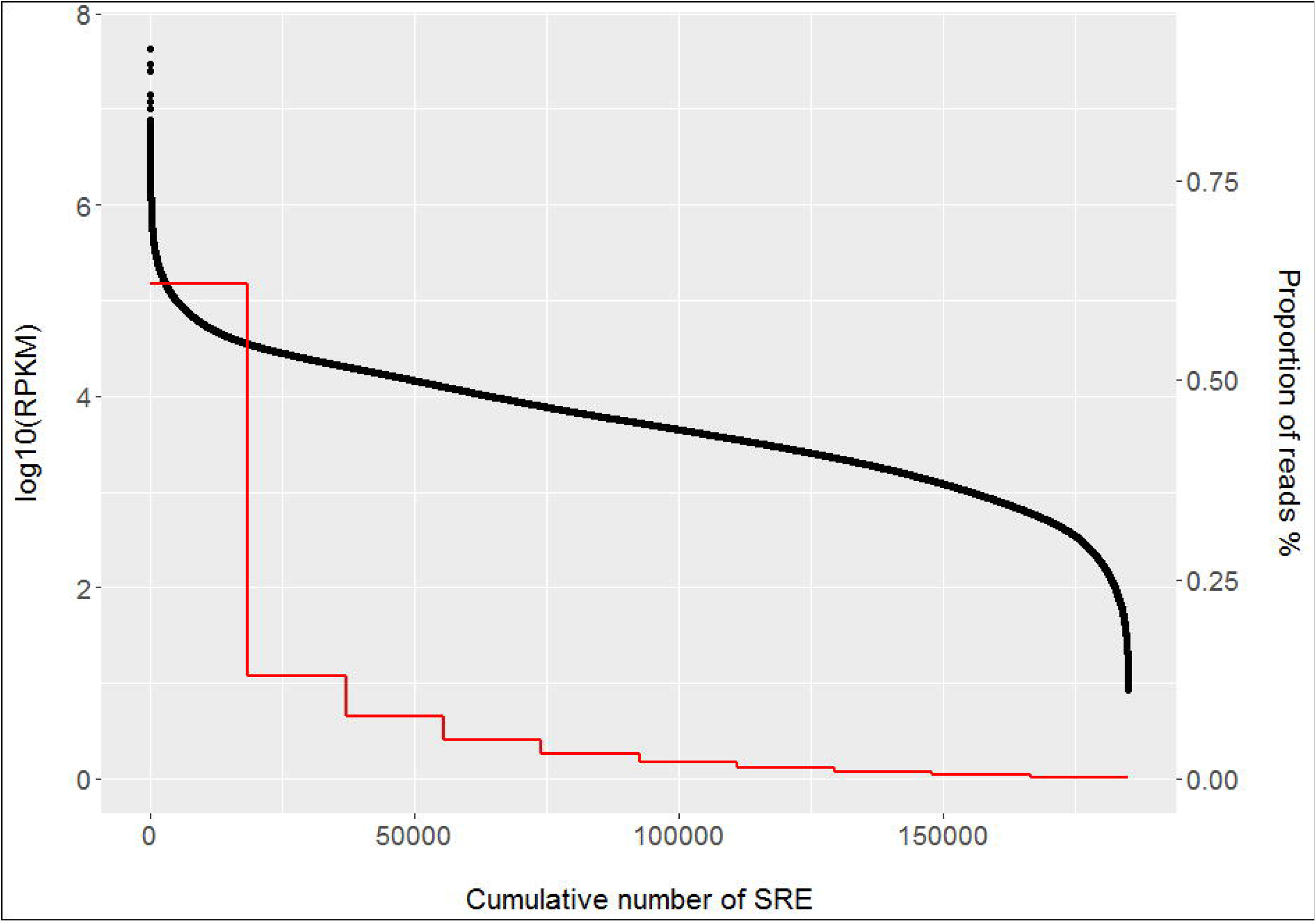
Cumulative abundance of the porcine SREs. The black dots indicate the log10 of the RNA abundance of each SRE. SREs are sorted in a decreasing order by their RNA abundance in the X axis. The red line represents the total number of SREs for each abundance decile group. The first decile of the most abundant SREs accounted for 65% of the total read abundance. RPKM: Reads Per Kilobase per Million mapped reads; SRE: Sperm RNA Element.

In order to dig further into the porcine sperm transcriptome, we investigated whether the orphan SRE syntenic regions in human and cattle included additional genes not annotated in pig. 1,505 and 1,313 SREs overlapped to syntenic regions in human and bovine, respectively. Forty five of the genes annotated within these regions were detected in both human and cattle (Supplementary File 3), including *CDYL*, a gene implicated in spermatid development and *ANXA3*, which protein levels in sperm have been found altered in men with poor semen compared to men with good sperm quality (Netherton et al., 2018). Ontology analysis of the 4,436 most abundant genes together with the 45 orphan SRE orthologs showed an enrichment of the cellular protein metabolic process (q-val: 2.7 x 10^-12^), macromolecular complex subunit organization (q-val: 2.1 x 10^-9^), sexual reproduction (q-val: 6.5 x 10^-8^), spermatogenesis (q-val: 1.2 x 10^-6^) and male gamete generation (q-val: 1.4 x 10^-6^), among others (Supplementary File 4). The transcripts detected in our study are concordant with previous results in human (Jodar et al., 2016) and bovine (Selvaraju et al., 2017) sperm and included genes related to fertilization (e.g. *HSPA1L* and *PRSS37*) or spermatogenesis (*ODF2* and *SPATA18*). The top 30 most abundant annotated protein coding SREs mapped to 27 genes (Table 1), 12 from mitochondrial origin (e.g. *COX1, COX2, ATP8, ATP6, COX3*), and 15 encoded in the nuclear genome (e.g. *PRM1, OAZ3, HSPB9, NDUFS4*). The abundance of mitochondrial genes reflects the high number of mitochondria typically contained in a spermatozoa cell to provide critical functions for the cell’s fertilizing ability including energy supply, regulation of molecular mechanisms involved in the development of the capacitation process, production of reactive oxygen species and calcium homeostasis (Rodriguez-Gil and Bonet, 2016). The 15 nuclear genes included members related to spermatogenesis, chromatin compaction and embryo development (Sendler et al., 2013; Selvaraju et al., 2017).

**Table 1.**
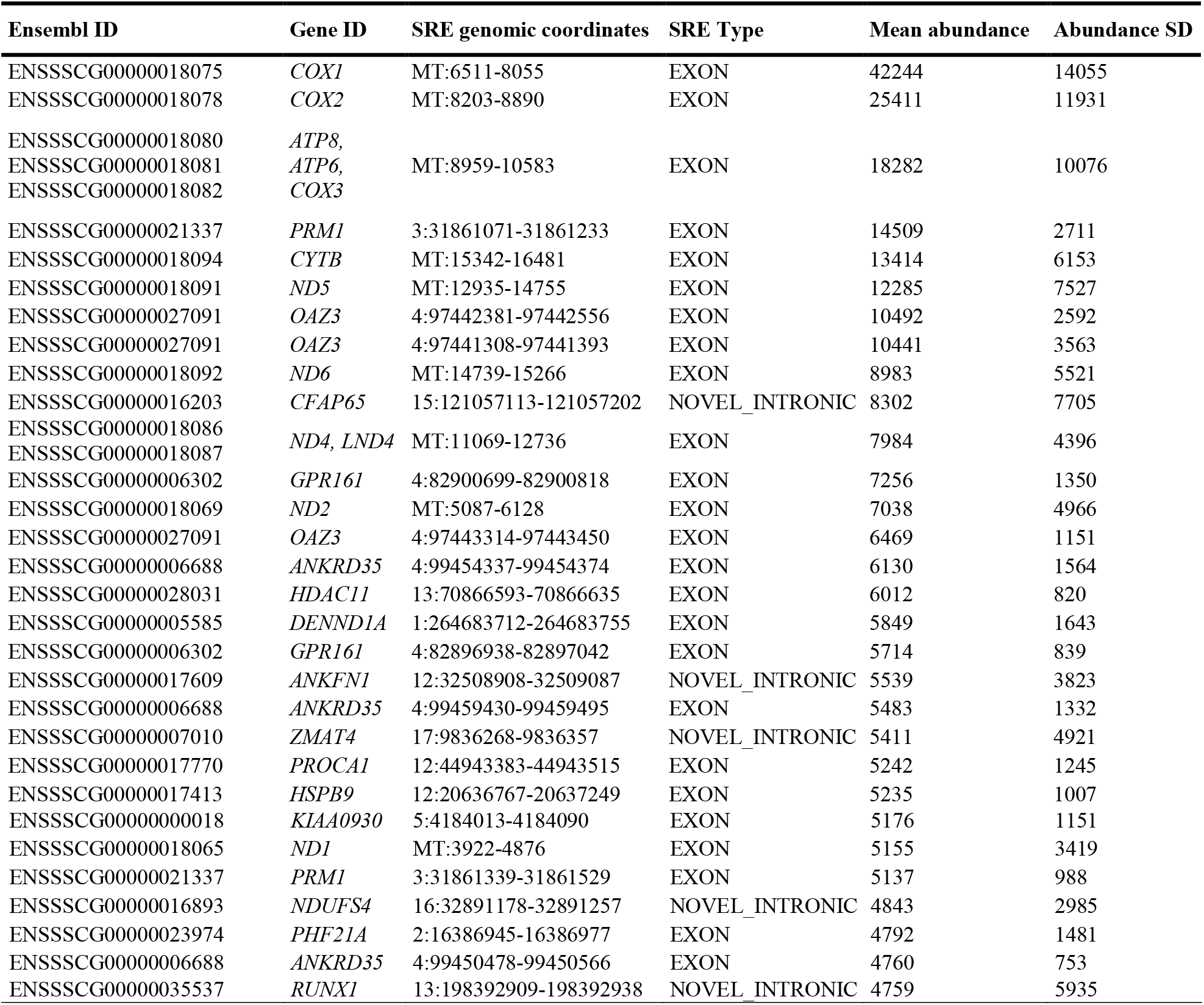
List of the 30 most abundant SREs in the porcine sperm. The most abundant SREs from protein coding genes included 12 mitochondrial and 15 nuclear genes. Some genes (e.g. *PRM1, OAZ3, ANKRD35*) presented more than one highly abundant SRE. SD: Standard Deviation; SRE: Sperm RNA Element. The SRE genomic coordinates are displayed in the format chromosome:start location–end location. Mean abundance and abundance SD are indicated in RPKM: Reads Per Kilobase per Million mapped reads.

### Total RNA-seq analysis: variance on the SRE abundance

We evaluated the transcripts that contained the 10% most abundant SREs across all samples and classified them as uniform (coefficient of variation or cv < 25%) or variable (cv > 75%). This identified 481 genes for which all their SREs were uniformly represented (cv < 25%) and 276 genes where each SRE was highly variable (cv > 75%). The list of 481 genes with constant abundance was enriched for several functions including the regulation of calcium, ATP generation and spermatid development and differentiation (Supplementary File 5). On the contrary, the highly variable genes were only enriched for the gene ontology term: single fertilization (zygote formation), which includes *SPMI, AQN-1* and *BSP1* among others (Supplementary File 5). This transcript variability is in general tolerated because it does not have severe phenotypic consequences. However, some of these transcripts may incur in a significant impact on semen quality and/or fertility and they could thus be biomarkers of the boar’s reproductive ability. Thus, it would be worth exploring the relationship between these genes and reproductive phenotypes in a larger study.

To further understand the functional relevance of the SRE abundance variability, we also searched for a potential relationship between this and the likely tissue of origin of the SRE. According to Jodar *et al*. sperm transcripts can be classified as testes-enriched, spermatozoa-enriched and seminal fluid-enriched (Jodar et al., 2016). Of the genes including the top decile SREs, 728, 381 and 448 were testes, spermatozoa and seminal fluid -enriched, respectively. We found a significant difference between the uniformity of the SRE abundance of the testes-enriched and the seminal fluid-enriched fractions (p-value: 3.6 x 10^-4^). The seminal fluid-enriched fraction was more variable. No difference in variability was found between the SREs of the sperm-enriched and the other fractions (p-values: 0.18 - 0.20). The lack of statistical difference between the sperm-enriched and the other fractions may be explained by two facts. On the one hand, a large proportion of the testes transcripts corresponds to cells belonging to the spermatogenic lineage. On the other hand, mature spermatozoa takes up seminal plasma RNAs via seminal exosomes (Vojtech et al., 2014; Jodar et al., 2016).

### Total RNA-seq analysis: Transcript Integrity

Sperm transcripts have been found to be highly fragmented in several mammalian species (Das et al., 2013; Sendler et al., 2013; Selvaraju et al., 2017; Gòdia et al., 2018a). We sought to investigate whether this fragmentation followed a programmatic pattern or perhaps was stochastic in the pig. For each annotated transcript, we calculated the abundance levels (in FPKM) and the TIN. In average, we found 31,287 protein coding transcripts with FPKM > 0 and TIN values > 0. Most transcripts (55%) were highly fragmented (TIN ≤ 25) whilst only 181 were almost intact (TIN > 75). Interestingly, the 10 samples showed similar TIN patterns across transcripts (Pearson correlation 0.72 - 0.93) (Supplementary File 6). The correlations between TIN and transcript length and transcript abundance were low (0.16 - 0.20 and 0.17 - 0.24, respectively) (Supplementary File 6). We then searched for gene ontology enrichment using the 10% most abundant transcripts within each TIN group. The highly fragmented group (TIN < 25) was enriched for genes related to negative regulation of JNK cascade (q-val = 1.2 x 10^-3^), spindle assembly (q-val = 5.6 x 10^-3^), and regulation of DNA repair (q-val = 4.5 x 10^-^), among others. These results are comparable to a previous study in human sperm (Sendler et al., 2013), where the most fragmented transcripts were not enriched for spermatogenesis or fertility functions. On the other hand, no significant pathways were found in the group of the top 10% most intact transcripts, possibly due to the low size of this group (18 transcripts), even though it contained genes related to spermatogenesis (*PRM1, OAZ3, ACSBG2*), sperm movement (*PRM3, SMCP*) or heat stress response (*HSPB9*) (Table 2). Remarkably, the six aforementioned genes were also within the most intact transcripts in human sperm (Sendler et al., 2013), thereby indicating conservation across species and their likely basic function in supporting sperm development and/or fecundity. Altogether, this indicates that the transcript fragmentation typically found in sperm may follow a programmatic basis and possible owe to relevant functions during spermatogenesis or upon fertilization.

**Table 2.** List of the 10% most abundant intact transcripts (TIN > 75) in the boar sperm. Transcript integrity was measured as TIN: Transcript Integrity Number; TIN mean: Average TIN. SD: Standard Deviation. * Gene symbol extracted from an orthologous sene species.

| Ensembl Transcript ID | Gene ID | TIN mean | TIN SD |
| --- | --- | --- | --- |
| ENSSSCT00000018955 | *ZNRF4* | 97.60 | 0.56 |
| ENSSSCT00000007842 | *TMEM239* | 93.54 | 2.54 |
| ENSSSCT00000019381 | *HSPB9* | 92.62 | 3.48 |
| ENSSSCT00000046661 | *UBL4B* | 91.93 | 2.24 |
| ENSSSCT00000001702 | *C6orf106* | 89.91 | 4.00 |
| ENSSSCT00000006503 | *SPATC1* | 86.40 | 2.61 |
| ENSSSCT00000030220 | *OAZ3* | 84.91 | 2.74 |
| ENSSSCT00000004015 | *AZIN2* | 83.92 | 3.15 |
| ENSSSCT00000049885 | *PRM3* | 83.56 | 2.27 |
| ENSSSCT00000029296 | *DBIL5\** | 83.21 | 3.71 |
| ENSSSCT00000014766 | *ZNRF4* | 82.36 | 2.23 |
| ENSSSCT00000048242 | *ACSBG2\** | 81.15 | 2.62 |
| ENSSSCT00000007224 | *SMCP* | 79.47 | 5.33 |
| ENSSSCT00000003898 | *KIF17* | 79.04 | 1.67 |
| ENSSSCT00000007327 | *ANKRD35* | 78.97 | 2.91 |
| ENSSSCT00000012714 | *DNAJB8* | 76.43 | 4.11 |
| ENSSSCT00000000746 | *TPI1* | 75.99 | 3.63 |
| ENSSSCT00000029974 | *PRM1* | 75.72 | 1.44 |

### Total RNA-seq analysis: *De novo* transcriptome assembly

We sought to further exploit the RNA-seq data by performing *de novo* assembly of the reads that did not map to the porcine genome. An average of 5.1 M unmapped reads per sample were used for the analysis (Supplementary File 1) and aligned into an average of 8,459 contigs per sample, with a median size (N50) of 259 bp (Supplementary File 7). These contigs were then contrasted by sequence homology against several databases and after filtering (see Methods), resulted in a list of 1,060 proteins from human, cattle, mouse, pig and other animal species with moderate to high RNA abundance (Supplementary File 8). Some of the proteins were detected in more than one species and accounted for a total of 768 unique genes (Supplementary File 9). The majority of these genes (739) were already present in the porcine annotation whilst 29 were classified as novel genes. From the annotated genes, 699 were also detected with our initial pipeline mapping the SREs to the porcine genome but 40 were only detected by this *de novo* assembly (Supplementary File 9). The reads that did not map to the genome but found a gene counterpart in the *de novo* analysis could have remained unmapped due to three main reasons. They could have either harbor more mismatches than the maximum allowed for the mapping algorithm, mapped within a repetitive element or, simply correspond to segments not assembled in the current version of the porcine genome. These three scenarios could involve full genes or just gene segments. The 40 known genes detected only by the de novo assembly together with the 29 potential novel genes did not cluster into any GO biological process. However, some of these genes have been associated to spermatogenesis or implicated in the sperm structure such as the sperm head or flagellum (e.g. *ACSBG2, HSF2BP, CCNYL1, KNL1* and *WBP2NL*). These results are in line with the recent study carried in humans by Perea and co-authors (Pertea et al., 2018) as already detailed in relation to the orphan SREs. Although the number of novel protein-coding genes represents a modest increase (29 genes), our de novo analysis yielded a much higher number (699) potentially novel splice variants.

### Total RNA-seq analysis: Repetitive Elements

REs are of particular interest as they comprise a high proportion of the porcine genome (approximately 40%) often related to genome instability (Bzymek and Lovett, 2001). Germline cells are very sensitive to the deleterious effects of active transposable elements. For example, the disruption of Long Interspersed Nuclear Element 1 (LINE1) retrotransposon silencing, the most abundant RE in the pig genome, can lead to spermatogenesis aberrations (Gòdia et al., 2018b) and embryo development arrest (Beraldi et al., 2006). Due to their relevance in spermatozoa, we annotated the RE segments that were transcribed in the pig sperm. A total of 4.6% of the mapped reads overlapped with REs, which is in line with previous data in murine sperm (Johnson et al., 2015), and accounted for 42.8 Mb of the swine genome. The most enriched RE classes included simple repeats (2.58% of the total mapped reads) which could potentially correspond to porcine nuclear matrix associated RNAs (Johnson et al., 2015). The second most abundant REs were the short Interspersed Nuclear Elements (SINEs) which accounted for 0.6% of the total read abundance. SINEs are transposable elements that can be hypo-methylated and can regulate male germ cell development, sperm packaging and embryo development (Schmid et al., 2001). In pigs, LINE1 accounts for 16.8% of the genome space and in our study, 0.19% of the mapped reads overlapped with LINE1 segments and spanned 25.5 Mb of the genome. This is nearly ten times less than in mice (1.89%) (Johnson et al., 2015) even though LINE1 is just slightly more ubiquitous in the murine genome (20%) (Waterston et al., 2002). While potentially interesting, these differences may arise due to yet unknown species-specific biological particularities or technical differences in the library preparation and/or bioinformatics methods used in both studies.

### Total RNA-seq analysis: long non-coding RNAs

lncRNAs are regulatory RNAs above 200 bp long implicated in a plethora of functions, including spermatogenesis and reproduction (Gòdia et al., 2018b). Sperm lncRNAs have been reported in human (Sendler et al., 2013), mice (Zhang et al., 2017) and cattle (Selvaraju et al., 2017). We identified 27 of the 361 lncRNA annotated in Ensembl v.91, and their RNA levels were clearly below their coding SREs counterparts (Supplementary File 10). The predicted cis-regulated target genes included *ZNF217*, which is a transcriptional repressor, *DYNLRB2* which encodes for a protein belonging to the dynein family of axoneme components related to sperm motility and *YIPF5*, which caused infertility in a knock-out fruitfly model (Yu et al., 2015). The annotation of lncRNAs in the swine genome remains remarkably poor and here we provide an initial catalogue that is still incomplete.

### Short RNA-seq analysis

On average, 6.6 M reads were obtained for each short RNA-seq library. A mean of 83% of these reads aligned to the queried porcine (*Sus scrofa*) databases (Supplementary File 1). A total of 34% of the aligned reads corresponded to sncRNAs, mainly piRNAs (37% of the sncRNA fraction), tRNAs (22.6%) and miRNAs (20.2%) (Figure 2 (A); Supplementary File 11). The remaining aligned reads (66%) mostly belonged to mitochondrial transfer and ribosomal RNAs (51%) but also to nuclear protein coding genes (Supplementary File 11).

**Figure 2.**
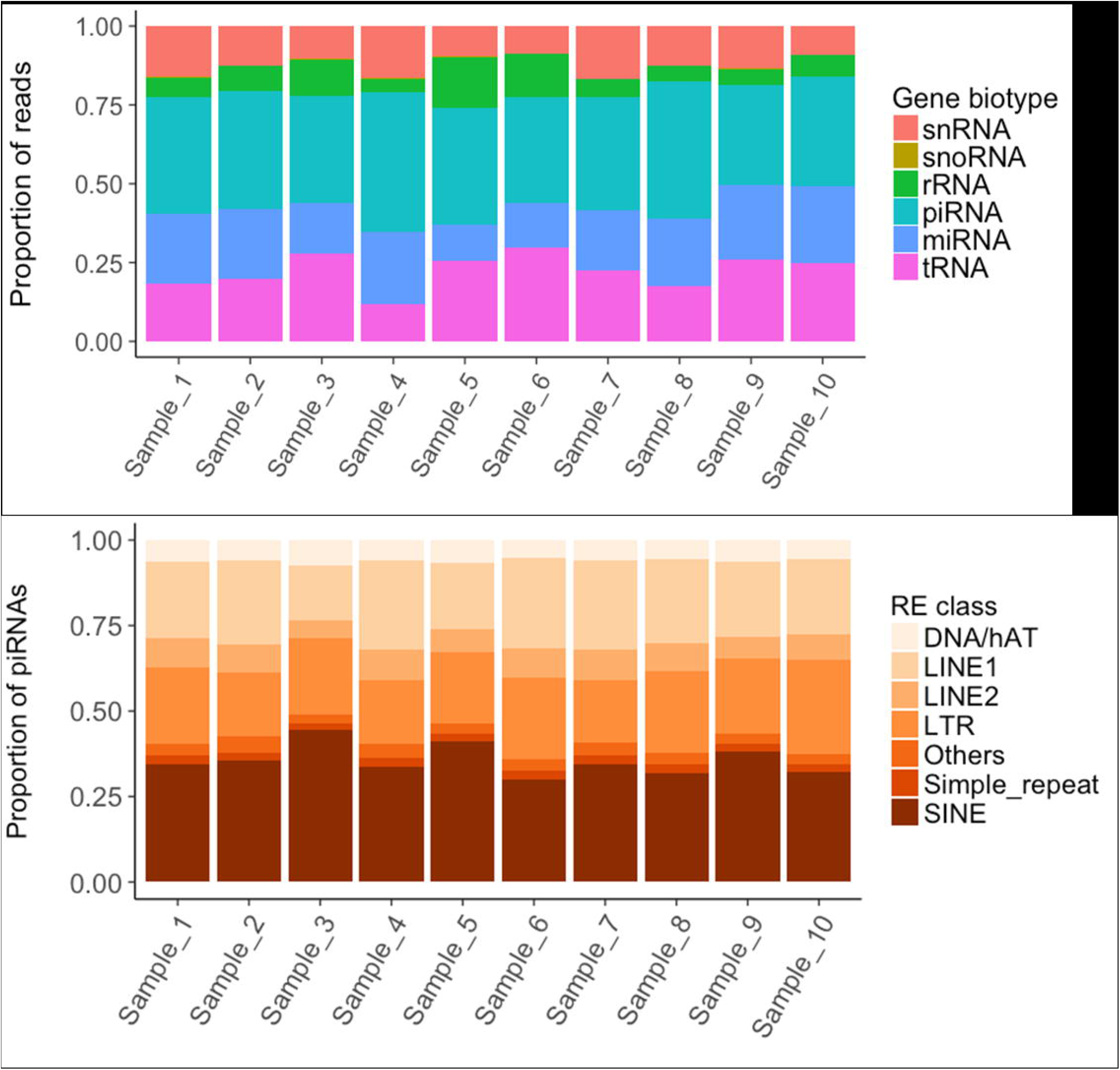
Read mapping distribution of the short non-coding RNA types and piRNA distribution within the Repetitive Element classes. **(A)**. Proportion of reads mapping to each short non-coding RNA type. **(B)**. Distribution within each Repetitive Element class of the piRNA cluster reads overlapping with Repetitive Elements.

The functional relevance of miRNAs, piRNAs and tRNAs in sperm biology and fertility (Krawetz et al., 2011; Gòdia et al., 2018b), (Sharma et al., 2016) is well known. miRNAs are a class of sncRNAs that have been found in multiple cell types and involved in a plethora of phenotypes and diseases. They post-transcriptionally repress the translation of target messenger RNAs (mRNAs) and can be ideal biomarkers for many traits including sperm quality and fertility. We detected 105 miRNAs (annotated in the pig) that were present in all the samples, with an average abundance that ranged from 4.6 to 13,192.2 counts per million (CPMs). Chen and colleagues (Chen et al., 2017a) detected a larger number of miRNAs (140), 75 of which were also detected in our experiment, in a RNA-seq study using 3 pooled pig sperm samples (Supplementary File 12). The reduced number of miRNAs described in our work is somewhat not surprising as we only considered those miRNAs that were present in the 10 samples. The inter-species comparison also indicates a degree of conservation in the miRNA composition of the mammalian sperm with about 70% of the miRNAs shared in cattle (Capra et al., 2017) and human (Pantano et al., 2015) (Supplementary File 12). These results suggest a conserved functional role of these miRNAs in mammals. The most abundant miRNAs in our study, miR-34c, miR-191, miR-30d, miR-10b and let-7a, among others (Supplementary File 13), are also highly abundant in cattle (Capra et al., 2017) and in human (Krawetz et al., 2011; Pantano et al., 2015) sperm. Some of these miRNAs have been linked to the male’s reproductive ability. For example, miR-34c is crucial for spermatogenesis (Yuan et al., 2015) and has been related to bull fertility (Fagerlind et al., 2015) and miR-191, miR-30d and miR-10b displayed altered levels in infertile human patients when compared to healthy controls (Salas-Huetos et al., 2015; Tian et al., 2017). We then assessed the coefficient of variation (cv) across the sperm samples to evaluate their abundance stability (Supplementary File 13). Interestingly, miRNAs showed large variability, 32% of them varied markedly (cv > 75%), including the highly abundant miR-34c, miR-30c-5p, miR-186 and miR-99a, with none showing low variability. As previously mentioned, exosome vesicles may also contribute in modulating the miRNA population of recipient cells. In fact, a recent study identified altered miRNA profiles in seminal plasma exosomes from azoospermic patients (Barcelo et al., 2018). We did not measure the pairwise correlation between the abundance of miRNAs and mRNAs because in their canonical function, miRNAs inhibit translation but have a small impact on the levels of the target mRNAs.

piRNAs are a class of 26-32 bp size sncRNAs that interact with Piwi proteins to contribute important functions to germline development, epigenetic regulation and the silencing of transposable elements (Gòdia et al., 2018b). We queried a public database of 501 piRNA clusters identified in pig testes (Rosenkranz, 2016), and found that 300 were represented in boar sperm and covered 5.03 Mb (0.20%) of the Scrofa10.2 genome assembly (Supplementary File 13). The RNA levels ranged between 3.2 and 5,242 CPMs and the cluster length between 5,077 and 114,717 bp. piRNA clusters tend to overlap with REs, in keeping with their role in genome inactivation and transposon regulation (Krawetz et al., 2011; Pantano et al., 2015; Gòdia et al., 2018b). In our work, 25% of the piRNA clusters co-localized with REs, most of which were SINEs (Figure 2 (B)). As piRNAs are tissue-specific and we queried a testes database (Rosenkranz, 2016), we also carried a *de novo* prediction of piRNA clusters with proTRAC using the remaining unaligned reads (average of 1.1 M reads) (Supplementary File 1). We identified 17 novel potential clusters of average abundance and length of 11.3 - 585 CPMs and 2,357 - 56,029 bp, respectively and as a whole, they covered 159.7 kb of the Sscrofa11.1 genome. Six of the novel clusters were present in the 10 samples and are thus considered of high confidence (Supplementary File 14).

tRNAs were the second most abundant class in porcine sperm, and their abundance is related to metabolic processes (Sharma et al., 2016). We identified 315 putative tRNAs from which 63% varied among samples (cv > 75%) (Supplementary File 13). Although the role of tRNAs in germ cells and in offspring health is uncertain, independent studies have shown that tRNA levels can be altered in response to certain manipulation of the paternal diet (Sharma et al., 2016; Gòdia et al., 2018b).

### Seasonal differences in the boar sperm transcriptome

A seasonal variation on semen quality and fertility has been observed in several animal species including the pig. During the warm summer months, as the scrotum is unable to thermo-regulate, spermatogenesis is negatively affected and the number of sperm cells and their motility tend to decrease alongside with an increase on morphological abnormalities (Zasiadczyk et al., 2015; Rodriguez et al., 2017). This effect on semen quality and also fertility (Suriyasomboon et al., 2006) has been related to heat stress. The molecular mechanisms underlying this phenomenon remain unclear although links to oxidative stress and the production of reactive oxidative species (ROS), with the consequent damage on sperm membrane integrity, DNA damage, apoptosis, autophagy and reduction of mitochondrial activity have been proposed (Durairajanayagam et al., 2015; Argenti et al., 2018). In a recent study, Argenti and co-authors (Argenti et al., 2018) identified increased superoxide dismutase anti-oxidant activity in the sperm of boars raised in sub-tropical Brazil in the summer months probably as a molecular attempt to reduce the presence of ROS and sperm damage (Argenti et al., 2018). Moreover, dietary strategies based on supplementary Zinc (Li et al., 2017) and l-arginine (Chen et al., 2018) have been related to a reduction of oxidative stress and improvement on the epididymal function and boar sperm quality in summer.

We compared the transcriptome (mRNA transcripts and miRNA) of the sperm samples collected in the summer months (May to August; N = 5) with those collected in winter (October to February; N = 5) in a temperate climate zone (latitude 42° N). The semen quality of the summer and winter groups was not significantly different although a trend was seen for sperm cell viability (p-val = 0.05), acrosome reaction (p-val = 0.09) and neck (p-val = 0.07) and tail (p-val = 0.08) morphological abnormalities. We detected 36 transcripts displaying a significant difference in abundance. Of these, two transcripts corresponded to the same gene and they were not taken into account due to concerns on the transcript allocation carried by the software. From, the 34 remaining transcripts, each from a different gene, 14 were up-regulated and 20 were down-regulated in the summer group (Table 3).

**Table 3.** Messenger RNA transcripts showing differential abundances in the summer versus the winter ejaculates. The list includes only these transcripts with q-val < 0.05 and log2 (FC) < −1.5 or >1.5. log2 (FC) > 0 indicate up-regulation in summer when compared to winter. Empty cells in the Gene ID column correspond to transcripts without gene symbol or description. FC: Fold-Change; FDR: False Discovery Rate.

| Transcript ID | Gene ID | Log2 (FC) | p-value | q-value (FDR) |
| --- | --- | --- | --- | --- |
| ENSSSCT00000058763 | *NSUN6* | -9.62 | 7.00E-16 | 4.35E-12 |
| ENSSSCT00000056639 | *ATG16L1* | -7.75 | 1.03E-08 | 2.14E-05 |
| ENSSSCT00000059752 | *EHBP1* | -7.49 | 7.95E-08 | 1.35E-04 |
| ENSSSCT00000059921 | *CENPC* | -6.55 | 8.56E-07 | 1.06E-03 |
| ENSSSCT00000012060 | *MTPAP* | -6.51 | 3.16E-07 | 4.21E-04 |
| ENSSSCT00000056608 | *SMARCA2* | -6.48 | 1.66E-05 | 1.15E-02 |
| ENSSSCT00000066205 | *CNOT3* | -6.22 | 1.75E-06 | 1.72E-03 |
| ENSSSCT00000014560 | *KIF18A* | -6.01 | 6.79E-05 | 3.62E-02 |
| ENSSSCT00000057538 | *ZNF24* | -5.88 | 4.01E-05 | 2.34E-02 |
| ENSSSCT00000018135 | *AOAH* | -5.47 | 2.54E-05 | 1.58E-02 |
| ENSSSCT00000015909 | *PSMD13* | -3.76 | 1.60E-05 | 1.15E-02 |
| ENSSSCT00000037719 | *STARD9* | -2.63 | 2.18E-05 | 1.45E-02 |
| ENSSSCT00000039055 | *CPEB3* | -2.32 | 4.98E-05 | 2.81E-02 |
| ENSSSCT00000039293 | *MED13L* | -2.13 | 1.62E-05 | 1.15E-02 |
| ENSSSCT00000043522 | *OSGIN1* | -1.75 | 9.49E-06 | 7.69E-03 |
| ENSSSCT00000012151 | *CUL2* | 1.66 | 5.66E-05 | 3.10E-02 |
| ENSSSCT00000001457 |  | 4.44 | 6.31E-14 | 2.94E-10 |
| ENSSSCT00000049515 | *ZMYND10* | 4.72 | 1.26E-06 | 1.31E-03 |
| ENSSSCT00000011652 | *TRUB1* | 4.93 | 2.33E-05 | 1.50E-02 |
| ENSSSCT00000049377 | *NUP58* | 5.14 | 9.56E-05 | 4.95E-02 |
| ENSSSCT00000007716 | *MCM8* | 5.15 | 1.68E-20 | 3.13E-16 |
| ENSSSCT00000035098 | *ERBIN* | 5.31 | 9.92E-06 | 7.71E-03 |
| ENSSSCT00000031111 | *ANKRD6* | 5.53 | 2.58E-06 | 2.40E-03 |
| ENSSSCT00000038311 | *MCPH1* | 5.65 | 4.31E-06 | 3.83E-03 |
| ENSSSCT00000018344 | *WDR70* | 5.72 | 1.04E-06 | 1.18E-03 |
| ENSSSCT00000037667 | *ASCC1* | 5.78 | 2.80E-05 | 1.69E-02 |
| ENSSSCT00000002542 | *FUT8* | 6.00 | 5.14E-06 | 4.36E-03 |
| ENSSSCT00000032033 | *TMEM230* | 6.01 | 2.08E-07 | 2.98E-04 |
| ENSSSCT00000050364 | *PDE3B* | 6.45 | 1.62E-07 | 2.52E-04 |
| ENSSSCT00000015769 | *FBXO38* | 6.48 | 3.51E-08 | 6.54E-05 |
| ENSSSCT00000043281 | *ZNF280D* | 6.49 | 1.08E-06 | 1.18E-03 |
| ENSSSCT00000064492 | *ZNF629* | 6.73 | 2.55E-09 | 6.80E-06 |
| ENSSSCT00000028805 | *ZNF583* | 7.34 | 6.81E-10 | 2.12E-06 |
| ENSSSCT00000030081 | *NMNAT1* | 7.50 | 1.59E-11 | 5.94E-08 |
| ENSSSCT00000039133 | *ATG16L1* | 7.76 | 4.16E-09 | 9.69E-06 |
| ENSSSCT00000038377 | *RUNDC3B* | 8.96 | 3.72E-16 | 3.47E-12 |

The largest difference in gene abundance between both seasonal groups (q-val = 3.13 x 10^-16^) corresponded to the minichromosome maintenance 8 homologous recombination repair factor *(MCM8)* gene (Table 3) which is a helicase related to the initiation of eukaryotic genome replication and may be associated with the length of the reproductive lifespan and menopause. *MCM8* plays a role in gametogenesis due to its essential functions in DNA damage repair via homologous recombination of DNA double strand breaks (Lutzmann et al., 2012). Another gene, the RUN Domain Containing 3B (*RUNDC3B*) has unknown functions but it contains a RUN domain that interacts with *RAP2*, a GTPase that has been linked to heat stress in plants (Figueroa-Yañez et al., 2016) and is related to male meiosis in mammals (Manterola et al., 2016). In keeping with *RAP2’s* function, a study in bulls found that spermatogonia undergoing meiosis during spermatogenesis were susceptible to heat stress (Rahman et al., 2018). This suggests that in mammals, spermatogonia exposed to heat stress, up-regulate the expression of *RUNDC3B* as a protective mechanism to ensure correct spermatogenesis and the production of normal spermatozoa. StAR Related Lipid Transfer Domain Containing 9 (*STARD9*), which was up-regulated in the summer group, is a lipid binding gene that has been related to asthenospermia in humans (Mao et al., 2011). Moreover, the paralog *STARD6* has been linked to spermatogenesis and spermatozoa quality (Mao et al., 2011). This is in keeping with the fact that the spermatozoon is very sensitive to oxidative damage for several reasons including the high amount of the peroxidation-prone unsaturated fatty acids that are present in its plasma membrane (Aitken and De Iuliis, 2010). Another gene that was found up-regulated in the summer group is the Oxidative Stress Induced Growth Inhibitor 1 gene *(OSGIN1). OSGIN1* has been related to autophagy and oxidative stress and its encoded protein regulates both cell death and apoptosis in the airway epithelium (Sukkar and Harris, 2017). Its expression is induced by DNA damage, which is one of the key sperm parameters that increase in the warm summer months (Perez-Crespo et al., 2008). Since this gene has also been identified in the sperm lineage, it could respond with a similar anti-oxidative role to heat stress in sperm.

The presence of RNA in ejaculated sperm in summer versus winter seasons has been previously interrogated using the microarray technology (Yang et al., 2010). In that study the authors identified 33 dysregulated transcripts, none of which was differentially abundant in our dataset. This lack of concordance between works could be due to both biological and technical reasons and is somewhat expected. First, the two studies interrogated different animal populations in different geographic locations. The study by Yang *et al*. (Yang et al., 2010) focused on Duroc boars breed in a sub-tropical region in Taiwan (25°N) whilst we screened Pietrain males from a sub-Mediterranean temperate climatic zone in Catalonia with warm summers and mildly cold winters (köppen classification Cfb; latitude 42°N). Moreover, we used a RNA-seq approach targeting the whole transcriptome whilst Yang and co-authors [21] employed a custom microarray interrogating only 708 target genes and by large, ignored the vast catalogue of annotated genes.

We also identified 5 miRNAs up- and 2 miRNAs down-regulated in summer (Table 4). This set included miR-34c, which was one of the most abundant miRNAs in our study, as well as in the sperm of other species, and was down-regulated in the summer samples. The RNA levels of miR-34c were also down-regulated in the sperm of men and mice exposed to severe early life stress events (Dickson et al., 2018), and in the testis of cynomolgus monkeys exposed to testicular hyperthermia (Sakurai et al., 2016), thus suggesting a link between the seasonality of semen quality and miR-34c. miR-1249, down-regulated in the summer group, was also found to be altered in the semen of bulls with moderate fertility (Fagerlind et al., 2015). Members of the miR-106 family were recently associated with oxidative stress in several tissues and cell types. For example, miR-106b targets the 12/15-Lipoxygenase enzymes, which are involved in the metabolism of fatty acids and oxidative stress in murine neurons (Wu et al., 2017). miR-106b has also been related to autophagy and cellular stress in intestinal epithelial HCT116 cells (Zhai et al., 2013). A study in cattle identified a single nucleotide polymorphisms in a miR-378 target site of the *INCENP* semen quality associated gene (Liu et al., 2016). In humans, miR-378 was found to also target the autophagy related protein 12 gene (*ATG12*) in cervical cancer (Tan et al., 2018). Finally, miR-221 was linked to autophagy in several tissues as well (Li et al., 2016; Qian et al., 2017) and was shown to regulate *SOD2*, which has key mitochondrial anti-oxidant functions in a murine model of ischemic skeletal muscle regeneration (Togliatto et al., 2013).

**Table 4.** List of the miRNAs showing distinct seasonal abundance. The list includes only these miRNAs with q-val < 0.05 and log2FC >1.5. FC: Fold-Change; FDR: False Discovery Rate.

| miRNA ID | Log2 (FC) | p-value | q-value (FDR) |
| --- | --- | --- | --- |
| ssc-miR-221-3p | -2.70 | 4.19E-05 | 1.54E-03 |
| ssc-miR-362 | -1.81 | 1.63E-03 | 2.18E-02 |
| ssc-miR-378 | -1.71 | 6.16E-03 | 4.94E-02 |
| ssc-miR-106a | -1.62 | 1.75E-05 | 1.29E-03 |
| ssc-miR-34c | -1.53 | 5.87E-04 | 9.59E-03 |
| ssc-miR-1306-5p | 1.68 | 1.81E-04 | 3.81E-03 |
| ssc-miR-1249 | 3.14 | 2.58E-08 | 3.79E-06 |

Our data, together with previous reports in swine, indicates that there is a molecular basis related to the well-reported decrease of semen quality and fertility in swine (Suriyasomboon et al., 2006; Zasiadczyk et al., 2015). These results should therefore be confirmed using additional animals and ideally, in a matched study where the winter and summer ejaculates come from the same boars. Nevertheless, our results are in keeping with previous data suggesting oxidative stress and autophagy as the key causes of the loss of semen quality in the warm summer periods.

## Conclusions

We have identified a rich and complex sperm transcriptome with known and novel coding RNAs, lncRNAs and sncRNAs that resembles the human, mouse and cattle counterparts. Their roles are mainly related to the regulation of spermatogenesis, fertility and early embryo development. These spermatozoal transcripts are fragmented, likely in a selective manner, consistently affecting some genes more than others across samples. This suggests that their fragmentation has a programmatic basis. Similarly, the variability of the transcript abundance between samples was transcript specific. This indepth transcriptome profile can be used a reference to identify RNA markers for semen quality and male fertility in pigs and in other animal species.

Interestingly, the levels of some transcripts changed between the summer and the winter ejaculates, most likely responding to heat stress, which would in turn, cause oxidative stress, sperm membrane and DNA damage and autophagy. Our data supports previous findings suggesting that feed supplementation can correct this seasonal effect and thus, opens the door to explore nutri-genomics research to improve semen quality and male fertility. The biological basis of these transcriptome changes needs to be further explored. In the recent years it has become evident that the ejaculate contains different sub-populations of sperm, each with specific roles upon ejaculation Thus, the changes in transcript abundances that we identified could reflect either similar variations on the transcript’s load in all spermatozoa cells or indicate alterations in the proportion of the sperm sub-populations each carrying their specific transcript profile. Discriminating both hypotheses could help defining the best strategies to mitigate this seasonal effect. Single-cell RNA-seq, a novel and powerful technology that still needs to be optimized in spermatozoa, could allow identifying the sperm sub-populations and their relevance for seasonality, semen quality and fertility. The trans-generational consequences in these transcript profiles are also worth the study. The altered RNA levels in sperm may perpetuate in the offspring’s ejaculate and have transgenerational phenotypic consequences. This should be also explored. In conclusion, our results pave the way to carrying future research to understand the molecular basis of semen quality seasonality in pigs, humans and other affected species.

## List of abbreviations

CPM: Counts per Million
CV: Coefficient of Variation
FPKM: Fragment per Kilobase per Million mapped reads
LINE1: Long Interspersed Nuclear Element 1
lncRNAs: Long non-coding RNAs
miRNAs: micro RNAs
NGS: Next Generation Sequencing
piRNAs: Piwi-interacting RNAs
RE: Repeat Element
RPKM: Reads Per Kilobase per Million mapped reads
SINE: Short Interspersed Nuclear Element
sncRNAs: small non-coding RNAs
SRE: Sperm RNA Element
TIN: Transcript Integrity Number
tRNAs: Transfer RNAs

## Declarations

### Ethics approval and consent to participate

The ejaculates obtained from pigs were privately owned for non-research purposes. The owners provided consent for the use of these samples for research. Specialized professionals at the farm obtained all the ejaculates following standard routine monitoring procedures and relevant guidelines. No animal experiment has been performed in the scope of this research.

### Availability of data and material

The datasets generated and/or analysed during the current study are available in the Gene Expression Omnibus repository, [PERSISTENT WEB LINK TO DATASETS]

### Competing interests

The authors declare that they have no competing interests

### Funding

This work was supported by the Spanish Ministry of Economy and Competitiveness (MINECO) under grant AGL2013-44978-R and grant AGL2017-86946-R and by the CERCA Programme/Generalitat de Catalunya. We thank the Agency for Management of University and Research Grants (AGAUR) of the Generalitat de Catalunya (Grant Number 2014 SGR 1528). We also acknowledge the support of the Spanish Ministry of Economy and Competitivity for the Center of Excellence Severo Ochoa 2016–2019 (Grant Number SEV-2015-0533) grant awarded to the Centre for Research in Agricultural Genomics (CRAG). MG acknowledges a PhD studentship from MINECO (Grant Number BES-2014-070560) and a Short-Stay fellowship from MINECO (EEBB-I-2017-12229) at S.A.K.’s laboratory. MS acknowledges a Rumble fellowship. Funds through Charlotte B. Failing Professorship to SAK are gratefully appreciated. ACl was recipient of a MINECO’s Ramon y Cajal research fellow (Grant Number RYC-2011-07763).

### Authors’ contributions

MG, AS and ACl conceived and designed the experiment; SB collected the samples and JERG carried the phenotypic analysis; MG performed sperm purifications and RNA extractions; ACa carried the qPCRs and their analysis; MG made the bioinformatics and statistic analysis; ME developed the SRE pipeline and provided bioinformatics support. MG analyzed the data, with special input from SAK and ACl. MG and ACl wrote the manuscript; all authors discussed the data and read and approved the contents of the manuscript.

## Supplementary Material

Supplementary_File_1.xlsx

RNA-seq quality and mapping statistics.

Average and Standard Deviation (SD) for the 10 boar sperm samples processed, including: amount of RNA extracted and several RNA-seq bioinformatics statistics for both total and small RNA-seq.

Supplementary_File_2.xl s

Distribution of the top decile most abundant SREs (Sperm RNA Elements) into SRE types and gene biotypes.

Number of SREs (within the top decile) for each SRE type (exonic, intronic, upstream/downstream 10 kb and orphan). Total non-redundant number of genes and their biotype for each SRE class.

Supplementary_File_3.xls

List of human and bovine genes identified by syntenic alignment of the orphan SREs. Orphan SRE genome coordinates were liftover to human and bovine coordinates, and the genes mapped in these regions were extracted. A total of 45 genes shared in both species were found. From these genes, 44 were already annotated in the *Sscrofa* Ensembl v.91 annotation. 17 of these genes were also detected by exonic, intronic and/or upstream / downstream 10 kb SREs. This suggests that orphan SREs could correspond to unannotated isoforms or to paralogous genes.

Supplementary_File_4.xls

Gene Ontology analysis of the genes including the top decile most abundant and the orphan SREs detected in the SRE pipeline.

GO biological process terms with significant Bonferroni corrected p-values (p-val < 0.05) and their associated genes.

Supplementary_File_5.xls

Gene Ontology analysis of the different SRE abundance variance groups.

GO biological process terms with significant Bonferroni corrected p-values (p-val < 0.05) and their associated genes.

Supplementary_File_6.xl s

Correlation between transcript integrity across samples, with transcript abundance and coding sequence length.

Correlation of the TIN (Transcripts Integrity Number) between samples, with the transcript abundance and with the coding sequence length of the transcripts.

This table shows the correlation of the TIN (Transcripts Integrity Number) between each pair of samples, the correlation of the TIN with the transcript average abundance in FPKM (Fragments per Kilobase per Million mapped reads) across the 10 samples, and the correlation of the TIN with the length of coding sequence of the transcripts.

Supplementary_File_7.xl s

Summary statistics of the *de novo* transcriptome assembly.

Summary statistics of the Trinity output based on the number of potential novel genes and transcripts, and size (in bp) of the contigs based on all transcripts isoforms or based only on the longest isoform for each potential gene.

Supplementary_File_8.xls

List of proteins identified by *de novo* analysis, with the species in which they were detected and transcript abundance.

*De novo* analysis of the unmapped reads resulted in 1,060 proteins which passed the quality control filters (see Methods). For each protein, we include the cognate species, the predicted RNA mean abundance in the 10 samples (in FPKM), the Standard Deviation (SD) of their RNA abundance and the gene ID symbol retrieved from Uniprot (https://www.uniprot.org/). FPKM: Fragments per Kilobase per Million mapped reads.

Supplementary_File_9.xl s

Non-redundant list of genes identified by *de novo* analysis.

768 potentially novel genes were identified from the unmapped reads. The gene symbol IDs were retrieved with Uniprot from the Trinity output protein names. These genes were detected in at least one species (detailed in column 2 of Supplementary File 8). The majority of these genes were annotated in the porcine Ensembl v.91 but 29 were identified as novel genes. 40 of the genes annotated in the porcine genome were not detected with the SREs pipeline which indicates that none of their cognate reads mapped to the genome even though these genes are annotaed.

Supplementary_File_10.xls

List of long non-coding RNAs detected in porcine sperm

Ensembl IDs of the lncRNAs identified in this study, their genome coordinates, average RNA abundance across the 10 samples and length. Most of the lncRNAs presented, as an average across all samples, low RNA abundances.

Supplementary_File_11.xls

Distribution of the short RNA-seq reads mapping to different RNA types.Proportion and Standard Deviation (SD) across the 10 samples.

Supplementary_File_12.xls

Concordance of miRNA identification between our dataset and other sperm RNA-seq studies.

Comparison of the miRNAs identified in our study with other sperm RNA-seq experiments in pig, in human and cattle.

Supplementary_File_13.xls

RNA abundance levels and coefficient of variation of miRNAs, tRNAs and piRNAs in the porcine sperm.

RNA abundance is measured in CPM (Counts Per Million) across the 10 samples. We only considered the miRNAS with > 0 CPMs in all the samples. The genomic coordinates of piRNAs refer to the Sscrofa10.2 built instead of Sscrofa11.1 as provided by the piRNAs cluster database [40].

Supplementary_File_14.xls

Novel piRNA clusters identified in the pig sperm RNA

We detected 17 potential clusters of piRNAs that were found in at least 3 of the 10 samples analysed in this study. Mean and Standard Deviation (SD) in CPM (Counts Per Million).

## References

Aitken, R.J., and De Iuliis, G.N. (2010). On the possible origins of DNA damage in human spermatozoa. Mol. Hum. Reprod. 16, 3–13. doi: 10.1093/molehr/gap059.

Argenti, L.E., Parmeggiani, B.S., Leipnitz, G., Weber, A., Pereira, G.R., and Bustamante-Filho, I.C. (2018). Effects of season on boar semen parameters and anti oxidant enzymes in the south subtropical region in Brazil. Andrologia. doi: 10.1111/and.12951.

Bao, W., Kojima, K.K., and Kohany, O. (2015). Repbase Update, a database of repetitive elements in eukaryotic genomes. Mobile DNA. 6, 11. doi: 10.1186/s13100-015-0041-9 (accessed August 2017).

Barcelo, M., Mata, A., Bassas, L., and Larriba, S. (2018). Exosomal microRNAs in seminal plasma are markers of the origin of azoospermia and can predict the presence of sperm in testicular tissue. Hum. Reprod. 33, 1087–1098. doi: 10.1093/humrep/dey072.

Beraldi, R., Pittoggi, C., Sciamanna, I., Mattei, E., and Spadafora, C. (2006). Expression of LINE-1 retroposons is essential for murine preimplantation development. Mol. Reprod. Dev. 73, 279–287. doi: 10.1002/mrd.20423.

Bindea, G., Mlecnik, B., Hackl, H., Charoentong, P., Tosolini, M., Kirilovsky, A., et al. (2009). ClueGO: a Cytoscape plug-in to decipher functionally grouped gene ontology and pathway annotation networks. Bioinformatics. 25, 1091–1093. doi: 10.1093/bioinformatics/btp101.

Bolger, A.M., Lohse, M., and Usadel, B. (2014). Trimmomatic: a flexible trimmer for Illumina sequence data. Bioinformatics. 30, 2114–2120. doi: 10.1093/bioinformatics/btu170.

Bruggmann, R., Jagannathan, V., and Braunschweig, M. (2013). In Search of Epigenetic Marks in Testes and Sperm Cells of Differentially Fed Boars. PLoS One. 8:e78691. doi: 10.1371/journal.pone.0078691.

Bzymek, M., and Lovett, S.T. (2001). Instability of repetitive DNA sequences: The role of replication in multiple mechanisms. Proc. Natl. Acad. Sci. U.S.A. 98, 8319–8325. doi: DOI 10.1073/pnas.111008398.

Capra, E., Turri, F., Lazzari, B., Cremonesi, P., Gliozzi, T.M., Fojadelli, I., et al. (2017). Small RNA sequencing of cryopreserved semen from single bull revealed altered miRNAs and piRNAs expression between High- and Low-motile sperm populations. BMC Genomics. 18, 14. doi: 10.1186/s12864-016-3394-7.

Chan, P.P., and Lowe, T.M. (2016). GtRNAdb 2.0: an expanded database of transfer RNA genes identified in complete and draft genomes. Nucleic Acids Res. 44, D184–D189. doi: 10.1093/nar/gkv1309.

Chen, C., Wu, H., Shen, D., Wang, S.S., Zhang, L., Wang, X.Y., et al. (2017a). Comparative profiling of small RNAs of pig seminal plasma and ejaculated and epididymal sperm. Reproduction. 153, 785–796. doi: 10.1530/Rep-17-0014.

Chen, J.Q., Li, Y.S., Li, Z.J., Lu, H.X., Zhu, P.Q., and Li, C.M. (2018). Dietary l-arginine supplementation improves semen quality and libido of boars under high ambient temperature. Animal. 12, 1611–1620. doi: 10.1017/S1751731117003147.

Chen, X.X., Che, D.X., Zhang, P.F., Li, X.L., Yuan, Q.Q., Liu, T.T., et al. (2017b). Profiling of miRNAs in porcine germ cells during spermatogenesis. Reproduction. 154, 789–798. doi: 10.1530/Rep-17-0441.

Das, P.J., McCarthy, F., Vishnoi, M., Paria, N., Gresham, C., Li, G., et al. (2013). Stallion Sperm Transcriptome Comprises Functionally Coherent Coding and Regulatory RNAs as Revealed by Microarray Analysis and RNA-seq. PLoS One. 8:e56535. doi: 10.1371/journal.pone.0056535.

Dickson, D.A., Paulus, J.K., Mensah, V., Lem, J., Saavedra-Rodriguez, L., Gentry, A., et al. (2018). Reduced levels of miRNAs 449 and 34 in sperm of mice and men exposed to early life stress. Transl. Psychiatry. 8, 101. doi: 10.1038/s41398-018-0146-2.

Durairajanayagam, D., Agarwal, A., and Ong, C. (2015). Causes, effects and molecular mechanisms of testicular heat stress. Reprod. Biomed. Online. 30, 14–27. doi: 10.1016/j.rbmo.2014.09.018.

Durinck, S., Spellman, P.T., Birney, E., and Huber, W. (2009). Mapping identifiers for the integration of genomic datasets with the R/Bioconductor package biomaRt. Nat. Protoc. 4, 1184–1191. doi: 10.1038/nprot.2009.97.

Fagerlind, M., Stalhammar, H., Olsson, B., and Klinga-Levan, K. (2015). Expression of miRNAs in Bull Spermatozoa Correlates with Fertility Rates. Reprod. Domest. Anim. 50, 587–594. doi: 10.1111/rda.12531.

Figueroa-Yañez, L., Pereira-Santana, A., Arroyo-Herrera, A., Rodriguez-Corona, U., Sanchez-Teyer, F., Espadas-Alcocer, J., et al. (2016). RAP2.4a Is Transported through the Phloem to Regulate Cold and Heat Tolerance in Papaya Tree (Carica papaya cv. Maradol): Implications for Protection Against Abiotic Stress. PLoS One. 11:e0165030. doi: 10.1371/journal.pone.0165030.

Gòdia, M., Mayer, F.Q., Nafissi, J., Castelló, A., Rodríguez-Gil, J.E., Sánchez, A., et al. (2018a). A technical assessment of the porcine ejaculated spermatozoa for a sperm-specific RNA-seq analysis. Syst. Biol. Reprod. Med. 64, 291–303. doi: 10.1080/19396368.2018.1464610.

Gòdia, M., Swanson, G., and Krawetz, S.A. (2018b). A History of Why Fathers’ RNA Matters. Biol. Reprod. 99, 147–159.

Grabherr, M.G., Haas, B.J., Yassour, M., Levin, J.Z., Thompson, D.A., Amit, I., et al. (2011). Full-length transcriptome assembly from RNA-Seq data without a reference genome. Nat. Biotechnol. 29, 644–652. doi: 10.1038/nbt.1883.

Jodar, M., Sendler, E., and Krawetz, S.A. (2016). The protein and transcript profiles of human semen. Cell Tissue Res. 363, 85–96. doi: 10.1007/s00441-015-2237-1.

Jodar, M., Sendler, E., Moskovtsev, S.I., Librach, C.L., Goodrich, R., Swanson, S., et al. (2015). Absence of sperm RNA elements correlates with idiopathic male infertility. Sci. Transl. Med. 7, 295re296. doi: 10.1126/scitranslmed.aab1287.

Johnson, G.D., Mackie, P., Jodar, M., Moskovtsev, S., and Krawetz, S.A. (2015). Chromatin and extracellular vesicle associated sperm RNAs. Nucleic Acids Res. 43, 6847–6859. doi: 10.1093/nar/gkv591.

Kim, D., Langmead, B., and Salzberg, S.L. (2015). HISAT: a fast spliced aligner with low memory requirements. Nat. Methods. 12, 357–360. doi: 10.1038/nmeth.3317.

Kozomara, A., and Griffiths-Jones, S. (2011). miRBase: integrating microRNA annotation and deep-sequencing data. Nucleic Acids Res. 39, D152–D157. doi: 10.1093/nar/gkq1027.

Krawetz, S.A., Kruger, A., Lalancette, C., Tagett, R., Anton, E., Draghici, S., et al. (2011). A survey of small RNAs in human sperm. Hum. Reprod. 26, 3401–3412. doi: 10.1093/humrep/der329.

Kuhn, R.M., Haussler, D., and Kent, W.J. (2013). The UCSC genome browser and associated tools. Brief. Bioinfomatics. 14, 144–161. doi: 10.1093/bib/bbs038.

Li, L., Wang, Z., Hu, X., Wan, T., Wu, H., Jiang, W., et al. (2016). Human aortic smooth muscle cell-derived exosomal miR-221/222 inhibits autophagy via a PTEN/Akt signaling pathway in human umbilical vein endothelial cells. Biochem. Biophys. Res. Commun. 479, 343–350. doi: 10.1016/j.bbrc.2016.09.078.

Li, Z., Li, Y., Zhou, X., Cao, Y., and Li, C. (2017). Preventive effects of supplemental dietary zinc on heat-induced damage in the epididymis of boars. J. Therm. Biol. 64, 58–66. doi: 10.1016/j.jtherbio.2017.01.002.

Liu, J., Sun, Y., Yang, C., Zhang, Y., Jiang, Q., Huang, J., et al. (2016). Functional SNPs of INCENP Affect Semen Quality by Alternative Splicing Mode and Binding Affinity with the Target Bta-miR-378 in Chinese Holstein Bulls. PLoS One. 11, e0162730. doi: 10.1371/journal.pone.0162730.

Love, M.I., Huber, W., and Anders, S. (2014). Moderated estimation of fold change and dispersion for RNA-seq data with DESeq2. Genome Biol. 15, 550. doi: 10.1186/s13059-014-0550-8.

Luo, Z.G., Liu, Y.K., Chen, L., Ellis, M., Li, M.Z., Wang, J.Y., et al. (2015). microRNA profiling in three main stages during porcine spermatogenesis. J. Assist. Reprod. Genet. 32, 451–460. doi: 10.1007/s10815-014-0406-x.

Lutzmann, M., Grey, C., Traver, S., Ganier, O., Maya-Mendoza, A., Ranisavljevic, N., et al. (2012). MCM8- and MCM9-deficient mice reveal gametogenesis defects and genome instability due to impaired homologous recombination. Mol Cell 47, 523–534. doi: 10.1016/j.molcel.2012.05.048.

Ma, X., Zhu, Y., Li, C., Xue, P., Zhao, Y., Chen, S., et al. (2014). Characterisation of Caenorhabditis elegans sperm transcriptome and proteome. BMC Genomics. 15, 168. doi: 10.1186/1471-2164-15-168.

Manterola, M., Sicinski, P., and Wolgemuth, D.J. (2016). E-type cyclins modulate telomere integrity in mammalian male meiosis. Chromosoma. 125, 253–264. doi: 10.1007/s00412-015-0564-3.

Mao, X.M., Xing, R.W., Jing, X.W., Zhou, Q.Z., Yu, Q.F., Guo, W.B., et al. (2011). [Differentially expressed genes in asthenospermia: a bioinformatics-based study]. Zhonghua Nan Ke Xue. 17, 694–698.

Marques, D.B.D., Lopes, M.S., Broekhuijse, M., Guimaraes, S.E.F., Knol, E.F., Bastiaansen, J.W.M., et al. (2017). Genetic parameters for semen quality and quantity traits in five pig lines. J. Anim. Sci. 95, 4251–4259. doi: 10.2527/jas2017.1683.

Martin, M. (2011). Cutadapt removes adapter sequences from high-throughput sequencing reads. EMBnet.journal. 17, 10–12. doi: 10.14806/ej.17.1.200

Netherton, J.K., Hetherington, L., Ogle, R.A., Velkov, T., and Baker, M.A. (2018). Proteomic analysis of good- and poor-quality human sperm demonstrates that several proteins are routinely aberrantly regulated. Biol. Reprod. 99, 395–408. doi: 10.1093/biolre/iox166.

OECD (2018). Meat consumption (indicator) [Online]. Available: https://www.oecd-ilibrary.org/content/data/fa290fd0-en [Accessed 15 July 2018].

Pantano, L., Jodar, M., Bak, M., Ballesca, J.L., Tommerup, N., Oliva, R., et al. (2015). The small RNA content of human sperm reveals pseudogene-derived piRNAs complementary to protein-coding genes. RNA. 21, 1085–1095. doi: 10.1261/rna.046482.114.

Perez-Crespo, M., Pintado, B., and Gutierrez-Adan, A. (2008). Scrotal heat stress effects on sperm viability, sperm DNA integrity, and the offspring sex ratio in mice. Mol. Reprod. Dev. 75, 40–47. doi: 10.1002/mrd.20759.

Pertea, M., Pertea, G.M., Antonescu, C.M., Chang, T.C., Mendell, J.T., and Salzberg, S.L. (2015). StringTie enables improved reconstruction of a transcriptome from RNA-seq reads. Nat. Biotechnol. 33, 290–295. doi: 10.1038/nbt.3122.

Pertea, M., Shumate, A., Pertea, G., Varabyou, A., Chang, Y.-C., Madugundu, A.K., et al. (2018). Thousands of large-scale RNA sequencing experiments yield a comprehensive new human gene list and reveal extensive transcriptional noise. bioRxiv. doi: 10.1101/332825.

Qian, L.B., Jiang, S.Z., Tang, X.Q., Zhang, J., Liang, Y.Q., Yu, H.T., et al. (2017). Exacerbation of diabetic cardiac hypertrophy in OVE26 mice by angiotensin II is associated with JNK/c-Jun/miR-221-mediated autophagy inhibition. Oncotarget. 8, 106661–106671. doi: 10.18632/oncotarget.21302.

Quinlan, A.R., and Hall, I.M. (2010). BEDTools: a flexible suite of utilities for comparing genomic features. Bioinformatics. 26, 841–842. doi: 10.1093/bioinformatics/btq033.

Rahman, M.B., Schellander, K., Luceno, N.L., and Van Soom, A. (2018). Heat stress responses in spermatozoa: Mechanisms and consequences for cattle fertility. Theriogenology. 113, 102–112. doi: 10.1016/j.theriogenology.2018.02.012.

Rodriguez-Gil, J.E., and Bonet, S. (2016). Current knowledge on boar sperm metabolism: Comparison with other mammalian species. Theriogenology. 85, 4–11. doi: 10.1016/j.theriogenology.2015.05.005.

Rodriguez, A.L., Van Soom, A., Arsenakis, I., and Maes, D. (2017). Boar management and semen handling factors affect the quality of boar extended semen. Porcine Health Manag. doi: 10.1186/s40813-017-0062-5.

Rosenkranz, D. (2016). piRNA cluster database: a web resource for piRNA producing loci. Nucleic Acids Res. 44, D223–D230. doi: 10.1093/nar/gkv1265.

Rosenkranz, D., and Zischler, H. (2012). proTRAC--a software for probabilistic piRNA cluster detection, visualization and analysis. BMC Bioinformatics. 13, 5. doi: 10.1186/1471-2105-13-5.

Rueda, A., Barturen, G., Lebron, R., Gomez-Martin, C., Alganza, A., Oliver, J.L., et al. (2015). sRNAtoolbox: an integrated collection of small RNA research tools. Nucleic Acids Res. 43, W467–W473. doi: 10.1093/nar/gkv555.

Sakurai, K., Mikamoto, K., Shirai, M., Iguchi, T., Ito, K., Takasaki, W., et al. (2016). MicroRNA profiles in a monkey testicular injury model induced by testicular hyperthermia. J. Appl. Toxicol. 36, 1614–1621. doi: 10.1002/jat.3326.

Salas-Huetos, A., Blanco, J., Vidal, F., Godo, A., Grossmann, M., Pons, M.C., et al. (2015). Spermatozoa from patients with seminal alterations exhibit a differential micro-ribonucleic acid profile. Fertil. Steril. 104, 591–601. doi: 10.1016/j.fertnstert.2015.06.015.

Schmid, C., Heng, H.H.Q., Rubin, C., Ye, C.J., and Krawetz, S.A. (2001). Sperm nuclear matrix association of the PRM1-> PRM2-> TNP2 domain is independent of Alu methylation. Mol. Hum. Reprod. 7, 903–911. doi: 10.1093/molehr/7.10.903.

Selvaraju, S., Parthipan, S., Somashekar, L., Kolte, A.P., Krishnan Binsila, B., Arangasamy, A., et al. (2017). Occurrence and functional significance of the transcriptome in bovine (Bos taurus) spermatozoa. Sci. Rep. 7, 42392. doi: 10.1038/srep42392.

Sendler, E., Johnson, G.D., Mao, S., Goodrich, R.J., Diamond, M.P., Hauser, R., et al. (2013). Stability, delivery and functions of human sperm RNAs at fertilization. Nucleic Acids Res. 41, 4104–4117. doi: 10.1093/nar/gkt132.

Sharma, U., Conine, C.C., Shea, J.M., Boskovic, A., Derr, A.G., Bing, X.Y., et al. (2016). Biogenesis and function of tRNA fragments during sperm maturation and fertilization in mammals. Science. 351, 391–396. doi: 10.1126/science.aad6780.

Sukkar, M.B., and Harris, J. (2017). Potential impact of oxidative stress induced growth inhibitor 1 (OSGIN1) on airway epithelial cell autophagy in chronic obstructive pulmonary disease (COPD). J. Thorac. Dis. 9, 4825–4827. doi: 10.21037/jtd.2017.10.153.

Suriyasomboon, A., Lundeheim, N., Kunavongkrit, A., and Einarsson, S. (2006). Effect of temperature and humidity on reproductive performance of crossbred sows in Thailand. Theriogenology. 65, 606–628. doi: 10.1016/j.theriogenology.2005.06.005.

Tan, D., Zhou, C., Han, S., Hou, X., Kang, S., and Zhang, Y. (2018). MicroRNA-378 enhances migration and invasion in cervical cancer by directly targeting autophagy-related protein 12. Mol. Med. Report. 17, 6319–6326. doi: 10.3892/mmr.2018.8645.

Tian, H., Li, Z.L., Peng, D., Bai, X.G., and Liang, W.B. (2017). Expression difference of miR-10b and miR-135b between the fertile and infertile semen samples (p). Forensic Sci. Int. Genet. Suppl. Ser. 6, E257–E259. doi: 10.1016/j.fsigss.2017.09.092.

Togliatto, G., Trombetta, A., Dentelli, P., Cotogni, P., Rosso, A., Tschop, M.H., et al. (2013). Unacylated ghrelin promotes skeletal muscle regeneration following hindlimb ischemia via SOD-2-mediated miR-221/222 expression. J. Am. Heart. Assoc. 2, e000376. doi: 10.1161/JAHA.113.000376.

Trudeau, V., and Sanford, L.M. (1986). Effect of season and social environment on testis size and semen quality of the adult Landrace boar. J. Anim. Sci. 63, 1211–1219.

Vojtech, L., Woo, S., Hughes, S., Levy, C., Ballweber, L., Sauteraud, R.P., et al. (2014). Exosomes in human semen carry a distinctive repertoire of small non-coding RNAs with potential regulatory functions. Nucleic Acids Res. 42, 7290–7304. doi: 10.1093/nar/gku347.

Wang, L., Wang, S., and Li, W. (2012). RSeQC: quality control of RNA-seq experiments. Bioinformatics. 28, 2184–2185. doi: 10.1093/bioinformatics/bts356.

Waterston, R.H., Lindblad-Toh, K., Birney, E., Rogers, J., Abril, J.F., Agarwal, P., et al. (2002). Initial sequencing and comparative analysis of the mouse genome. Nature. 420, 520–562. doi: 10.1038/nature01262.

Wettemann, R.P., Wells, M.E., Omtvedt, I.T., Pope, C.E., and Turman, E.J. (1976). Influence of elevated ambient temperature on reproductive performance of boars. J. Anim. Sci. 42, 664–669.

Wood, D.E., and Salzberg, S.L. (2014). Kraken: ultrafast metagenomic sequence classification using exact alignments. Genome Biol. 15, R46. doi: 10.1186/gb-2014-15-3-r46.

Wu, Y., Xu, D., Zhu, X., Yang, G., and Ren, M. (2017). MiR-106a Associated with Diabetic Peripheral Neuropathy Through the Regulation of 12/15-LOX-meidiated Oxidative/Nitrative Stress. Curr. Neurovasc. Res. 14, 117–124. doi: 10.2174/1567202614666170404115912.

Yang, C.C., Lin, Y.S., Hsu, C.C., Tsai, M.H., Wu, S.C., and Cheng, W.T. (2010). Seasonal effect on sperm messenger RNA profile of domestic swine (Sus Scrofa). Anim. Reprod. Sci. 119, 76–84. doi: 10.1016/j.anireprosci.2009.12.002.

Yang, C.C., Lin, Y.S., Hsu, C.C., Wu, S.C., Lin, E.C., and Cheng, W.T.K. (2009). Identification and sequencing of remnant messenger RNAs found in domestic swine (Sus scrofa) fresh ejaculated spermatozoa. Anim. Reprod. Sci. 113, 143–155. doi: 10.1016/j.anireprosci.2008.08.012.

Yu, J., Wu, H., Wen, Y., Liu, Y.J., Zhou, T., Ni, B.X., et al. (2015). Identification of seven genes essential for male fertility through a genome-wide association study of non-obstructive azoospermia and RNA interference-mediated large-scale functional screening in Drosophila. Hum. Mol. Genet. 24, 1493–1503. doi: 10.1093/hmg/ddu557.

Yuan, S., Tang, C., Zhang, Y., Wu, J., Bao, J., Zheng, H., et al. (2015). mir-34b/c and mir-449a/b/c are required for spermatogenesis, but not for the first cleavage division in mice. Biol. Open. 4, 212–223. doi: 10.1242/bio.201410959.

Zasiadczyk, L., Fraser, L., Kordan, W., and Wasilewska, K. (2015). Individual and seasonal variations in the quality of fractionated boar ejaculates. Theriogenology. 83, 1287–1303. doi: 10.1016/j.theriogenology.2015.01.015.

Zhai, Z., Wu, F., Chuang, A.Y., and Kwon, J.H. (2013). miR-106b fine tunes ATG16L1 expression and autophagic activity in intestinal epithelial HCT116 cells. Inflamm. Bowel Dis. 19, 2295–2301. doi: 10.1097/MIB.0b013e31829e71cf.

Zhang, X., Gao, F., Fu, J., Zhang, P., Wang, Y., and Zeng, X. (2017). Systematic identification and characterization of long non-coding RNAs in mouse mature sperm. PLoS One. 12, e0173402. doi: 10.1371/journal.pone.0173402.

